# Deep-sea siliceous sponges harbor distinct and functionally diverse microbiomes

**DOI:** 10.64898/2026.03.04.709506

**Authors:** Katherine R. Lane, Kirstin S. Meyer-Kaiser, Adena B. Collens, Camille V. Leal, Allen G. Collins, Santiago Herrera, Colleen M. Hansel

## Abstract

Sponges, phylum Porifera, are long-lived and basal-branching metazoans that play important roles in ocean biogeochemistry and host diverse microbial communities. Siliceous sponges form a major clade of the Porifera, yet their microbiome is not well-characterized, particularly in the deep ocean. Here, we used shotgun metagenomics to investigate the composition of the microbial communities of 13 siliceous sponges collected from four sites near Puerto Rico from depths ranging from 400-1900 meters. Nine of the sponges in this study are from five sponge family taxa that have not previously been sequenced using shotgun metagenomics. We assembled a total of 176 metagenome-assembled-genomes from 20 bacterial and 1 archaeal phyla. Ammonia-oxidizing archaea (AOA) *Nitrosopumilaceae* dominated most siliceous sponge microbial communities and was strikingly the sole symbiont associated with one sponge (*Farrea*). Overall, microbiome diversity was relatively low across siliceous sponges, except for a Phloeodictyidae, which was likely a high microbial abundance (HMA) sponge. Our results suggest that host sponge phylogeny may shape microbial community structure, with limited evidence for environmental influence. The sponge-associated microbial communities contained genetic capabilities for diverse metabolic functions, particularly contributing to the carbon, nitrogen, and sulfur cycles. In addition to the AOA, evidence of potential for microbial autotrophy was found through the presence of genes for RuBisCO, methanotrophy, and ATP citrate lyase. These results reveal both conserved relationships and metabolic flexibility across siliceous sponge lineages, suggesting unique evolutionary dynamics and demonstrating the importance of microbial metabolism to sponge host health and nutrient cycling in the oligotrophic deep ocean.

**Importance:** Marine sponges, emerging ∼600 million years ago, have close relationships with microorganisms, but the microbiome of deep-sea siliceous sponges is not well understood. Siliceous sponges play essential roles in deep-sea ecosystems by providing habitats for other metazoans and mediating carbon, nitrogen, and sulfur cycling, yet they remain some of the least studied sponges. By shotgun sequencing DNA from 13 siliceous sponges collected near Puerto Rico, this study found that host sponge phylogeny influences microbial community composition and structure. Ammonia-oxidizing archaea dominated the microbial communities associated with marine sponges, likely playing key roles in utilizing metabolic byproducts and supporting host health. Other microbes also contributed to nutrient cycling and contained the potential to fix carbon, suggesting metabolic flexibility which may benefit sponge hosts in low-resource environments. These findings emphasize the ecological importance of siliceous sponge-microbe symbioses and contribute to our understanding of the drivers shaping their structure and function.

## 1.0 Introduction

Sponges, phylum Porifera, arose almost 600 million years ago (C.-W. Li et al. 1998; Love et al. 2009; Shawar et al. 2025) and are one of the earliest diverging extant animals in the tree of life (Schultz et al. 2023), playing important roles in the ocean’s past and present ecosystems. Siliceous sponges consist of silica-based species in classes Hexactinellida, Demospongiae, and Homoscleromorpha. Hexactinellid sponge reefs dominated margins and rocky outcrops in the Tethys Sea during the Jurassic (Bayer et al. 2020; Sally P. Leys 2003; S. P. Leys et al. 2007). Previously thought to be extinct, hexactinellid bioherm reefs were discovered in the 1990s off the coast of British Columbia (BC, Canada) (Conway et al. 1991; Krautter et al. 2001). These reefs form structure and habitat for complex and diverse ecosystems, creating hotspots of biodiversity and silica sinks (Maldonado et al. 2017).

The ecological and evolutionary success of marine sponges may in part be due to their symbiotic relationships with microbes, which are known to play crucial roles in metabolism and immunity (Wilkins et al. 2019; Woodhams et al. 2020). The sponge microbiome is known to contribute to nitrogen cycling, vitamin production, and novel secondary metabolite production, supporting both host health and the broader ecosystem (Glasl et al. 2024; de Goeij et al. 2013; Woodhouse et al. 2013).

The drivers that shape the microbiome acquisition and composition are not fully understood in Porifera (Lurgi et al. 2019; Thomas et al. 2016), particularly in deep ocean sponges (Busch et al. 2022b). Microbe-host interactions are shaped by the host organism and/or the environment. Acquisition of the microbiome may occur through vertical transmission (maternally to offspring), horizontal transmission (environmental), or a combination of the two (reviewed in Wilkins et al. 2019; Koskella et al. 2017). Previous research using operational taxonomic units (OTUs) to characterize microbial composition has shown patterns of host specificity in sponge-associated microbiome composition and structure (Busch et al. 2022a; Jackson et al. 2013; Reveillaud et al. 2014; Steinert et al. 2020). Furthermore, researchers have found highly conserved relationships between archaeal symbionts and hexactinellids using OTU sequencing (Garritano et al. 2023), suggesting the importance of host phylogeny in siliceous sponge-microbe symbioses.

Porifera are filter-feeders, pumping large amounts of water each day with recorded rates of up to 6.48 Ls^-1^kg^-1^ (Weisz et al. 2008) and thereby transforming the surrounding seawater chemistry. Often referred to as the “sponge loop,” sponge-driven circulation of seawater couples the benthic and pelagic ocean and recycles nutrients, thus playing a key role in the productivity of coral reefs and the deep ocean (de Goeij et al. 2013; Soest et al. 2012). The relative contributions of sponge host metabolism vs. associated microorganisms’ metabolism in carbon, nitrogen, and sulfur cycling remain ambiguous (Fiore et al. 2013; Hanz et al. 2022; Maldonado et al. 2021).

Despite their important roles in biogeochemical cycling in marine ecosystems, deep-sea siliceous sponges remain some of the least-studied sponge groups because they are mostly restricted to the deep ocean (Busch et al. 2022a; Wörheide et al. 2012; Ramirez-Llodra et al. 2010). Yet, because of their long evolutionary history and long life spans (i.e., hundreds to thousands of years (Jochum et al. 2012; Schultz et al. 2023)), hexactinellids and siliceous demosponges present a unique opportunity to consider the evolutionary pressures that shape host-microbe interactions.

Recent technological advances are greatly expanding our knowledge of deep-sea siliceous sponges. In the last two decades, remotely operated vehicles (ROVs) have enabled research on the ecology and physiology of deep marine organisms, including siliceous sponges (Maldonado et al. 2021). High-throughput sequencing technologies and accompanying advances in computational analysis have supported rapid progress in the genomics of siliceous sponges (Schultz et al. 2023; Steffen et al. 2023) and their associated microbiomes (Bayer et al. 2020; Garritano et al. 2024; Santini et al. 2023; Wei et al. 2023).

This study expands recent progress in sponge microbiome research by characterizing the microbiome of siliceous sponges collected between 400 to 1900 meter depths in the Caribbean. Of the thirteen sponges included in this study, nine are from five sponge families that have not been sequenced genetically before. Here, we combine sponge genomic and morphological methods to identify host sponge phylogeny at a high resolution with metagenomic techniques to investigate the associated microbiome composition and metabolic potential. We investigate the potential roles of environment (i.e., collection location) and sponge phylogeny in shaping microbiome structure and composition, providing insight into the relationships between host and microbes of these unique and understudied organisms.

## 2.0 Methods

### 2.1 Sample Collection

Siliceous sponge specimens were collected during the *R/V Falkor (too)* research cruise FKt230417 along the coast of Puerto Rico, April 17 – May 6, 2023. Sampling was opportunistic, depending on the availability of sponges in the deep-sea, and space availability in the multi-chamber BioBox containers filled with ambient seawater during the dive. High-resolution 4K video and images of each sampled sponge were recorded using ROV SuBastian (Figure 1: Sponge Plate). Specimens were collected using ROV SuBastian’s soft grip manipulators, to keep the sponges intact, including their base (peduncle). When possible, sponges were sampled with their attached substrate so that specimens were kept fully intact; in soft sediments, sponges were gently pulled upwards to extract the peduncle. After sampling, sponges were placed in the BioBox on ROV SuBastian’s front basket and brought to the surface at the end of the dive. Depth, oxygen concentration, and salinity at the time and location of sample collection were measured by a sensor package on ROV SuBastian (Figure 2: Depths and Map). Metadata associated with all specimens in this study are detailed in Supplemental Table 1.

**Figure 1.**
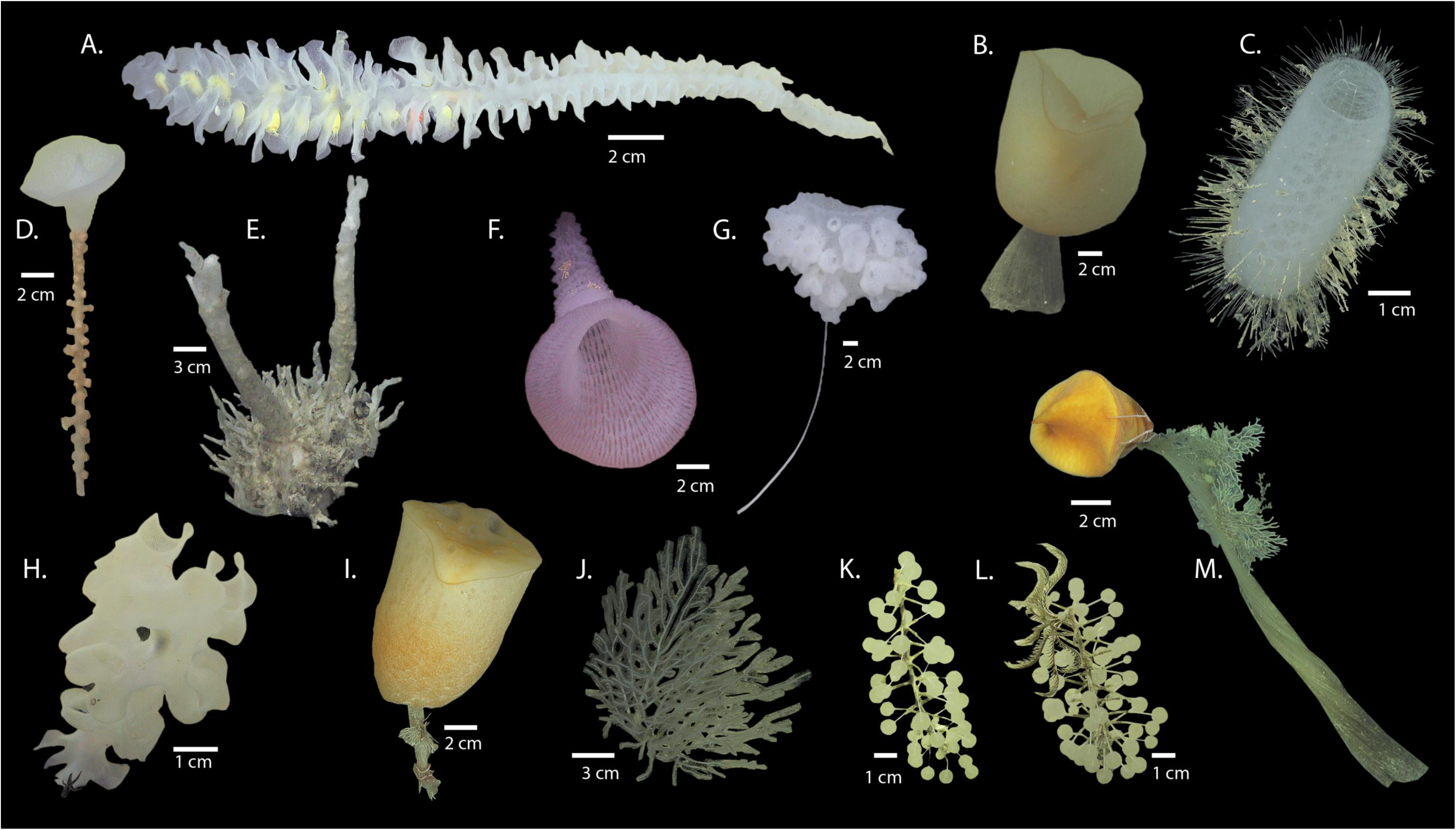
Siliceous sponge specimens *in situ.* Siliceous sponge specimens sampled along the Puerto Rican coast during R/V *Falkor (too)* research cruise FKt230417, April 17 – May 6, 2023, photographed using the high-resolution video camera (4K) on ROV SuBastian. **A:** Euretidae, *Pleurochorium* SH18746; **B:** Hyalonematidae, *Hyalonema* SH18914; **C:** Euplectellidae, *Euplectella* SH18731; **D:** Hyalonematidae, *Hyalonema* SH18879; **E:** Phloeodictyidae SH18796; **F:** Euretidae, *Lefroyella* SH18833; **G:** Euplectellidae, *Hyalostylus* SH18781; **H:** Farreidae, *Farrea* SH18656; **I:** Hyalonematidae, *Hyalonema* SH18724; **J:** Cladorhizidae, *Cladorhiza* SH18875; **K:** Cladorhizidae, *Chondrocladia* SH18747; **L:** Cladorhizidae, *Chondrocladia* SH18725; **M:** Hyalonematidae, *Hyalonema* SH18898.

**Figure 2.**
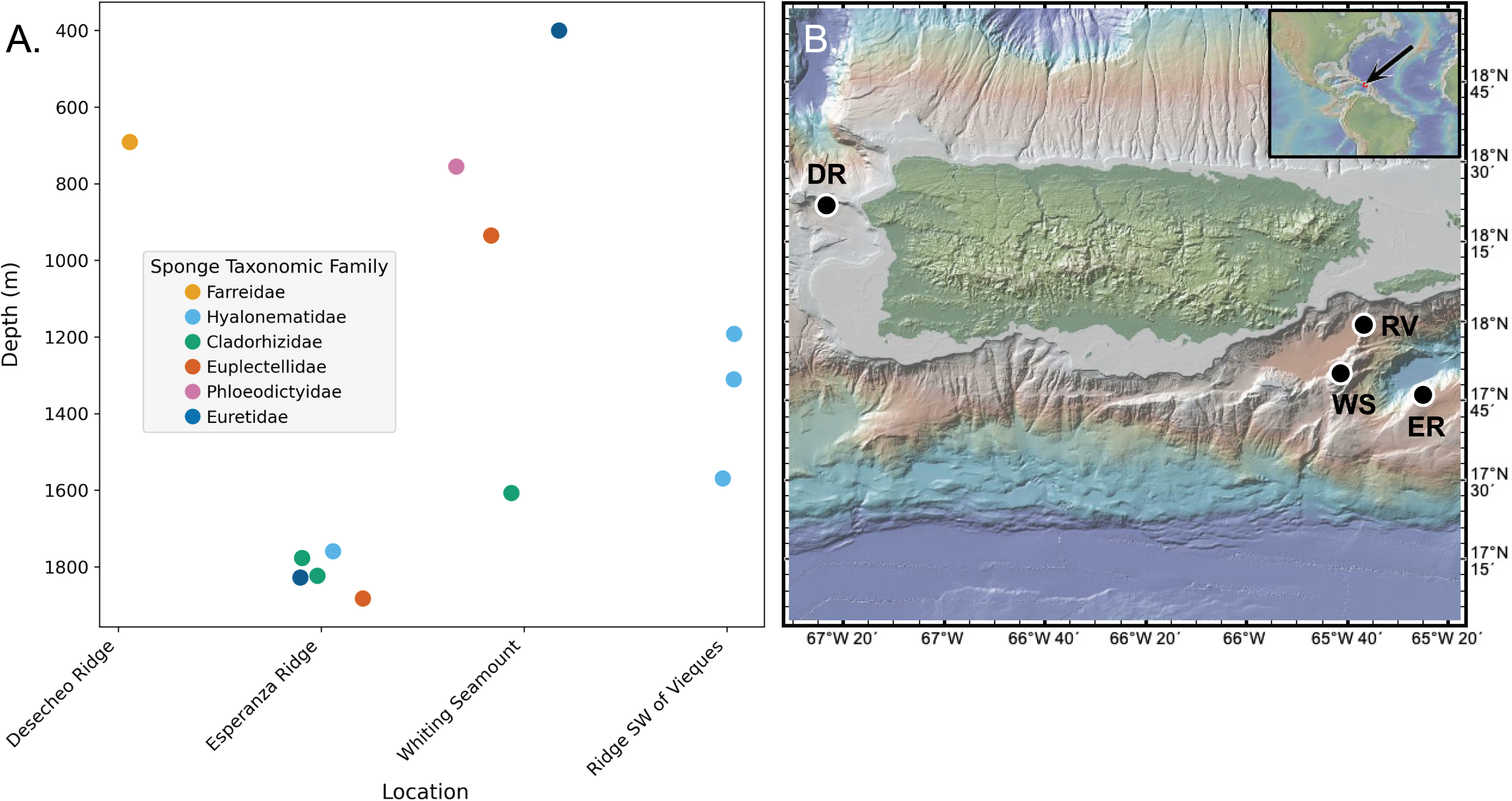
Sampling depths and locations. A: Distribution of sponge specimens in this study across sites and depths. Sponge specimens colored by taxonomic family collected at each site and plotted by depth (Cladorhizidae = Green, Euretidae = Dark Blue, Euplectellidae = Orange, Farreidae = Yellow, Hyalonematidae = Light Blue, Phloeodictyidae = Pink). B: Locations of the four dive sites where specimens were collected (Desecheo Ridge = DR, Esperanza Ridge = ER, Whiting Seamount = WS, Ridge SW of Vieques = RV).

A total of 13 siliceous sponges were collected: 9 hexactinellids and 4 demosponges. Following recovery, specimens were photographed with a ruler for scale. Sponges were rinsed thoroughly with sterile seawater and subsectioned with a sterilized ceramic knife upon a sterilized Teflon sheet. For each sponge, 2-3 half-centimeter diameter sections were sub-sampled from the upper part of the sponge (surface and internal to ensure representative sampling of the sponge microbiome). Subsections of siliceous sponges were flash-frozen in sterile WhirlPak bags and stored at –80°C for later metagenomic sequencing of the microbiome. A sub-sample of tissue (0.5 cm diameter) from each sample was preserved in ethanol and deposited in the collections of the National Museum of Natural History (NMNH), Smithsonian Institution, Washington DC and used for taxonomic characterization and phylogenetic analysis.

### 2.2 DNA Extraction and Sequencing

DNA was extracted from frozen sponge tissue samples using the DNA Powersoil Kit (Qiagen) with the following modifications: first vortex was 6 minutes, elution was performed twice (half of the recommended volume each time), and solution C6 was incubated for 3 minutes on the column. DNA was quantified using the Qubit 2.0 fluorometer (Life Technologies). Due to low biomass and low DNA yield, pooled DNA (2-3 extractions for each sponge) from each specimen was shipped on dry ice to Roy J. Carver Biotechnology Center at the University of Illinois at Urbana-Champaign for library prep and sequencing. The libraries for metagenome sequencing were prepared from genomic DNA extracts with the Illumina DNA Prep Kit (Illumina). Due to low yield DNA, sponges Cladorhizidae *Chondrocladia* SH18725 and SH18747; Euplectellidae *Hyalostylus* SH18781; Euplectellidae *Euplectella* SH18731; Euretidae *Lefroyella* SH18833; and Hyalonematidae *Hyalonema* SH18879 had 5 additional PCR cycles of amplification. Sponges Farreidae *Farrea* SH18656 and Euretidae *Pleurochorium* SH18746 had 8 additional PCR cycles of amplification. Additional PCR cycles can potentially introduce amplification biases in relative abundance estimates in the microbiome (Silverman et al. 2021); however, the low DNA yields of these low-biomass samples necessitated their use. DNA was sequenced with Illumina NovaSeq S4 paired end (PE) reads 150bp with sample read counts ranging from 226,490,692 reads (specimen SH18746) to 339,372,306 reads (specimen SH18875) (Supplemental Table 2). Raw reads are available in the National Center for Biotechnology Information (NCBI) Sequence Read Archive (SRA) BioProject Accession: PRJNA1355197.

### 2.3 Computational Analysis of Microbiome

Metagenomic raw reads were trimmed, cleaned, and quality controlled with BBDuk v38.84 (Bushnell et al. 2017) and FastQC v0.11.9 (Andrews 2010). Sample SH18898 was sequenced twice: the top and bottom of the sponge, and these sequencing reads were pooled. Metagenomes were assembled with Megahit v1.2.9 (D. Li et al. 2015) with default parameters. Contigs shorter than 1000bp were removed from downstream analyses, and assemblies were binned into metagenome-assembled genomes (MAGs) with Metabat2 v2.17 (Kang et al. 2019) with default parameters. Reads were mapped to the MAGs with BWA-MEM (H. Li and Durbin 2009) and Samtools (H. Li et al. 2009). Bins were quality controlled with Checkm2 v1.02 (Completeness > 70%, Contamination < 10%) (Parks et al. 2015), coverage for each bin was calculated with CoverM v0.7.0 (Aroney et al. 2025) and normalized by total number of mapped reads per sample so that relative abundances summed to 1. Taxonomy was assigned with GTDB-Tk v207 (Chaumeil et al. 2020). Genes were predicted with Prodigal v2.6.3 ‘-metà mode (Hyatt et al. 2010). Functional annotation was performed with METABOLIC-C v4.0 (Zhou et al. 2022), a Hidden Markov Model based tool that uses Kyoto Encyclopedia of Genes and Genomes (KEGG) database (Ogata et al. 1999), Carbohydrate-Active enZymes database (Lombard et al. 2014), and a set of custom HMMS (Johnson et al. 2010). Computational analysis was performed on Woods Hole Oceanographic Institution’s High Performance Computing Cluster Poseidon.

### 2.4 Sponge Taxonomic and Phylogenetic Assignment

*In situ* and shipboard images were used to assess external morphology. Subsampled tissues were used to prepare spicule slides and skeleton sections following (Stoddart and Johannes 1978) to characterize internal anatomy. The morphological characters of each specimen were compared with poriferan taxonomic literature. The taxonomic identification was verified and refined by conducting a phylogenetic analysis using the full-length SSU-rRNA gene tree with sponge reference sequences from SILVA (v. 138.2) and (Dohrmann et al. 2023). Separately from the microbiome computational analysis (Section 2.3), raw reads of the 13 metagenomes were quality-trimmed with fastp (Chen et al. 2018). Full-length SSU rRNAs were then assembled with phyloFlash (Gruber-Vodicka et al. 2020). Within phyloFlash, reads were mapped to SSU rRNA reference sequences in the SILVA 138.2 database (Quast et al. 2013; Yilmaz et al. 2014). Then, only those reads which had mapped to the references were assembled de-novo with SPAdes (Bankevich et al. 2012). For specimen SH18724, the full-length SSU rRNA was not assembled, so reads were re-mapped using Bowtie2 (Langmead and Salzberg 2012) to references, and a consensus sequence was taken from the highest coverage and identity mapping. Specimen SH18914 did not contain sufficient reads mapping to Hexactinellid SSU rRNA for phylogenetic resolution, thus it was excluded from the phylogenetic tree and included in further analyses using only the morphological identification.

To determine the phylogenetic position, the SSU rRNA from the 12 specimens were aligned with the most recent phylogeny of Hexactinellida SSU rRNA using MAFFT v7.490 (Katoh and Standley 2013) with the auto alignment algorithm and default parameters, from which a phylogenetic hypothesis was inferred using maximum likelihood in RAxML v8.2.11 assuming the GTR-GAMMA model of nucleotide evolution, with node support assessed using bootstrap resampling (Stamatakis 2014). A phylogenetic tree using only the 12 SSU-rRNA genes from sponges in this study was re-aligned, visualized, and formatted using iTOL (Letunic and Bork 2021) (Figure 3A, Supplemental File 1).

**Figure 3.**
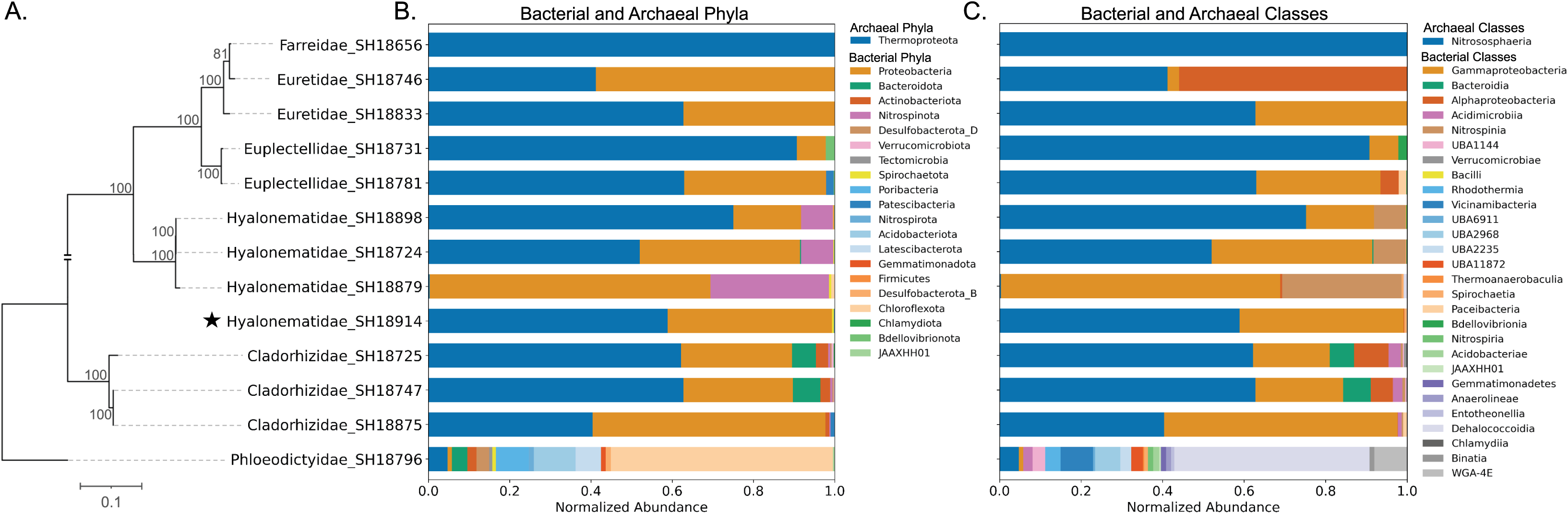
Sponge host phylogeny and associated microbial community composition. Phylogenetic tree (A) based on sponge SSU rRNA shows evolutionary relationships among sponge specimens. Specimen Hyalonematidae SH18914 (indicated with star), did not have sufficient Hexactinellid SSU rRNA data for inclusion in the tree, it was placed amongst the other specimens based on morphological taxonomic identification. Normalized abundance of Bacteria and Archaea Phylum (B) and Class (C) according to GTDB-Tk taxonomic assignment in each sponge specimen obtained in this study.

### 2.5 Statistical Analysis

Statistical analyses of community composition were performed in R (v. 4.5.1) using packages *vegan* (Oksanen 2017) and *pairwiseAdonis* (Martinez Arbizu 2020). Each sponge specimen was treated as a single sample, and analyzed data consisted of normalized abundance (coverage) data of each specimen’s microbial taxa at the phylum level. Multivariate statistical analyses were conducted based on a Bray-Curtis dissimilarity matrix using taxonomic levels of microbial phyla and sponge family. PERMANOVA was used to test for significant differences in microbial community composition based on sponge taxonomic family and collection location. Due to low sample sizes and limited residual degrees of freedom, two-way PERMANOVA was used to test the interaction term between sponge family collection location with the ‘by = “margin”’ parameter, which uses Type III sum of squares approach. PERMANOVA was used to test for differences within sponge Families with specimens from multiple locations. Pairwise post hoc PERMANOVA between sponge families were performed to compare group differences. Non-metric multidimensional scaling (nMDS) was used to visualize patterns in microbial community composition. A similarity percentages test (SIMPER) was used to identify the primary microbial phyla contributing to each sponge family’s dissimilarity. Simpson evenness (Simpson 1949) was calculated as a summary metric for microbiome evenness between sponge families, and a two-way crossed ANOVA was used to test for significant differences in Simpson evenness based on sponge taxonomic family and collection location. A Levene’s test was used to test homogeneity of variance of Simpson evenness values. Tukey HSD post hoc test was used to identify differences between groups in significant main effects and interactions of Simpson evenness values. For families with specimens collected from multiple locations, one-way ANOVAs were used to test for within-family differences in Simpson evenness across locations. Figures were made using the *ggplot* package in R (Wickham 2016) and Python. Code used for data analysis is available on Github: (https://github.com/kate-lane/siliceous-sponge-microbes).

## 3.0 Results

### 3.1 Sponge Taxonomy

All 13 siliceous sponge specimens in this study were identified to the family level using genomic and morphological analysis (Table 1; Figure 3A). Of the hexactinallids, four families were represented in sampling: Euretidae (n = 2), Euplectellidae (n = 2), Hyalonematidae (n = 4), and Farreidae (n = 1). Of the Demosponges, two families were sampled: Cladorhizidae (n = 3) and Phloeodictyidae (n = 1). Of the sponges in this study, the Phloeodictyidae was the only likely high microbial abundance (HMA) sponge (Moitinho-Silva, Steinert, et al. 2017) compared to the others which were categorized as low microbial abundance (LMA) sponges (Figure 3).

**Table 1.** Sponge taxonomy and accession numbers.

| Table 1 \| Sponge Taxonomy |  |  |  |  |  |  |  |
| --- | --- | --- | --- | --- | --- | --- | --- |
| Sam<br>ple | Class | Order | Family | Genus | Morphological<br>Analysis<br>(Smithsonian) | Genomic<br>Analysis | USNM<br>(Smithsonian<br>Collections) |
| SH18<br>656 | Hexactin<br>ellida | Sceptrul<br>ophora | Farreidae | *Farrea* | Yes | Yes | 1689167 |
| SH18<br>724 | Hexactin<br>ellida | Amphidis<br>cosida | Hyalone<br>matidae | *Hyalone*<br>*ma* | Yes | Yes | 1752114 |
| SH18<br>725 | Demosp<br>ongiae | Poecilosc<br>lerida | Cladorhiz<br>idae | *Chondro*<br>*cladia* | Yes | Yes | 1721463 |
| SH18<br>731 | Hexactin<br>ellida | Lyssacino<br>sida | Euplectell<br>idae | *Euplecte*<br>*lla* | No | Yes | N/A |
| SH18<br>746 | Hexactin<br>ellida | Sceptrul<br>ophora | Euretidae | *Pleuroch*<br>*orium* | Yes | Yes | 1757515 |
| SH18<br>747 | Demosp<br>ongiae | Poecilosc<br>lerida | Cladorhiz<br>idae | *Chondro*<br>*cladia* | Yes | Yes | 1757516 |
| SH18<br>781 | Hexactin<br>ellida | Lyssacino<br>sida | Euplectell<br>idae | *Hyalosty*<br>*lus* | Yes | Yes | 1757523 |
| SH18<br>796 | Demosp<br>ongiae | Oceanoa<br>pia | Phloeodi<br>ctyidae | - | Yes | Yes | 1689402 |
| SH18<br>833 | Hexactin<br>ellida | Sceptrul<br>ophora | Euretidae | *Lefroyell*<br>*a* | Yes | Yes | 1752122 |
| SH18<br>875 | Demosp<br>ongiae | Poecilosc<br>lerida | Cladorhiz<br>idae | *Cladorhi*<br>*za* | Yes | Yes | 1752123 |
| SH18<br>879 | Hexactin<br>ellida | Amphidis<br>cosida | Hyalone<br>matidae | *Hyalone*<br>*ma* | Yes | Yes | 1750689 |
| SH18<br>898 | Hexactin<br>ellida | Amphidis<br>cosida | Hyalone<br>matidae | *Hyalone*<br>*ma* | Yes | Yes | 1752124 |
| SH18 | Hexactin | Amphidis | Hyalone | *Hyalone* | Yes | Yes | 1752120 |
| 914 | ellida | cosida | matidae | *ma* |  |  |  |

The phylogenetic tree overall demonstrated close evolutionary relationships within specimens from the same families. We identified strong support for all clades (bootstrap = 100%), except for a clade including Farreidae_SH18656 and Euretidae_SH18746 (bootstrap = 81%). The low support for this clade and limited resolution between families Euretidae and Farreidae shown here is likely due to the close genetic relationship between Euretidae and Farreidae, for which there is evidence that Euretidae may be paraphyletic with respect to Farreidae (Reiswig and Dohrmann 2014; Dohrmann et al. 2017, 2023). The Phloeodictyidae sponge was the longest-branching from the other sponges in this study, which was likely explained by faster mutation rates in Haplosclerida (Niamh E. Redmond et al. 2011; N. E. Redmond et al. 2013) rather than evolutionary distance from the other sponges (Figure 3A).

### 3.2 Microbial Community Composition and Abundance

Metagenome-assembled genomes (MAGs) in our 13 sponge specimens included 21 microbial phyla (20 Bacteria and 1 Archaea), and 29 microbial classes (28 Bacteria and 1 Archaea) (Figure 3).

Microbial communities in our sponge samples were dominated by ammonia-oxidizing archaea (AOA) of class *Nitrosopheara*: family *Nitrosopumilaceae* (detailed in Supplemental Table 3). The microbiome of Farreidae SH18656 contained only two strains of archaeal family *Nitrosopumilaceae*. One of the Hyalonematidae SH18879 did not contain a high abundance of AOA, but contained *Nitrospinota* (nitrite-oxidizing bacteria) in higher abundance than other sponge specimens. In total, three of the *Hyalonema* contained *Nitrospinota* in higher abundances, and the two Cladorhizidae (SH18725 and SH18747) also contained *Nitrospinota* but in lower abundance.

The second-most abundant phylum was *Proteobacteria*, which was also present in all the sponge microbiomes, represented with classes *Gammaproteobacteria* and *Alphaproteobacteria*.

Two of the Cladorhizidae, (SH18725 and SH18747) were highly similar in their microbiome. They also contained bacterial phylum *Bacteriodota* in higher abundance than the other sponges.

Two of the sponges (Euplectellidae SH18781 and Cladorhizidae SH18875) contained *Patescibacteria*, a bacterial phylum from the Candidate Phyla Radiation. Euplectellidae SH18731 was the only sponge to contain phylum *Bdellovibrionata*.

### 3.3 Controls on Sponge Microbiome Composition

Despite the limited sample sizes per sponge family and site in this study, we conducted statistical analyses to investigate trends within the dataset. It is noted, however, that more robust statistical interpretations would require larger sample sizes to fully validate the results here.

A visual representation of microbial community composition showed that sponges clustered loosely by family (color) but not location (shape) (Figure 4). Note that Phloeodictyidae SH18796 was excluded from the presented nMDS because as the only likely HMA specimen it was an outlier in its microbial community (see Supplemental Figure 1). Two PERMANOVA tests showed that microbial community composition was significantly different based on sponge family (F_5,7_ = 5.1101, *p =* 0.01) but not collection location (F_3,9_ = 1.1752, *p =* 0.237). The interaction term between sponge family and collection location was significant (2-way crossed PERMANOVA, by = margin, F_2,3_ = 7.3422, *p =* 0.031). However, of the four sponge families with specimens from multiple collection sites, none of them showed significant statistical differences across sites (post hoc PERMANOVAs, *p* > 0.05). Family Cladorhizidae showed site-specific clustering in microbial community composition in the NMDS ordination (Fig. 4). The two Cladorhizidae that clustered together (specimens K and L) were both in genus *Chondrocladia*, while specimen J belonged to genus *Cladorhiza*. Thus, apparent clustering by location in this dataset may have actually been driven by phylogeny at lower taxonomic levels.

**Figure 4.**
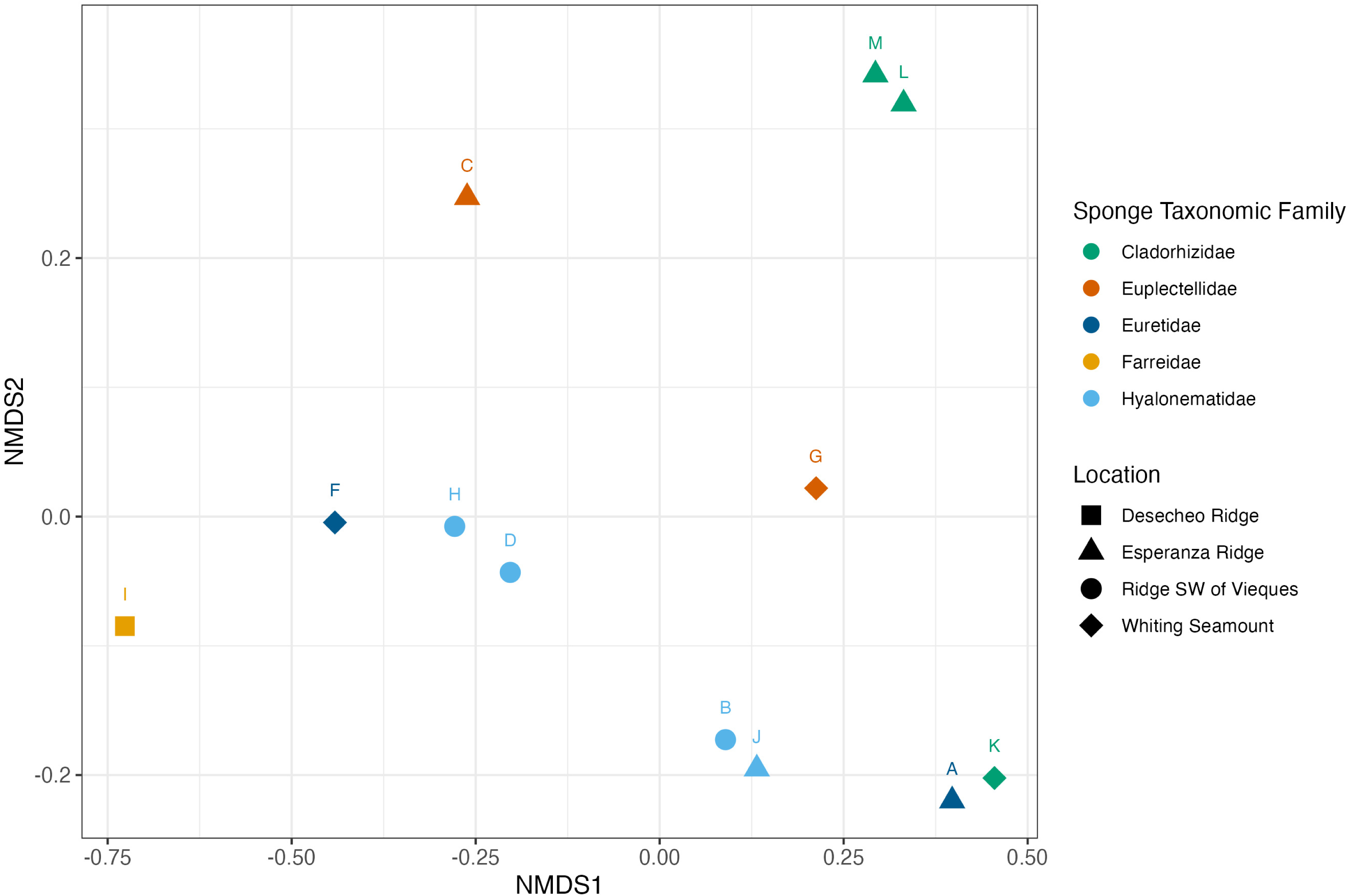
nMDS ordination of microbial community composition across siliceous sponges. Non-metric multidimensional scaling plot (nMDS) based on Bray Curtis similarities of sponge microbiome phyla composition and abundance. Closely clustered symbols correspond to samples with similar Bacteria and Archaeal phyla composition and abundance. Sponge family taxa are indicated by color. Locations of specimens are indicated by shape (Desecheo Ridge = Square, Esperanza Ridge = Triangle, Ridge SW of Vieques = Circle, Whiting Seamount = Diamond.) Letters adjacent to symbols indicate individual specimens as labelled in Figure 1: Sponge Plate.

A pairwise *post hoc* PERMANOVA test showed that only one pair of sponge families had significantly different microbial community composition: Hyalonematidae vs Cladorhizidae (uncorrected *p =* 0.015), but this difference was not significant after multiple testing correction (adjusted *p* = 0.434). None of the other pairs of sponge families in the pairwise post hoc test had significantly different microbial community composition, likely due to the low numbers of replicates.

Simpson evenness differed significantly between sponge families (2-way crossed ANOVA, F_5,3_= 24.17, *p =* 0.0125) but not collection locations (F_2,3_= 3.65, *p =* 0.157) (Figure 5). There was a significant interaction between sponge family and location (two-way crossed ANOVA, F_2,3_= 12.48, *p =* 0.0351). Post hoc tests (Tukey’s HSD) revealed that Cladorhizidae at Esperanza Ridge and Phloeodictyidae at Whiting Seamount had significantly higher evenness than Farreidae at Desecheo Ridge (*p =* 0.0363) and (*p* = 0.0349), respectively. No other pairwise differences in Simpson evenness between families across locations were significant, including within-family statistical tests of Simpson evenness across locations of the four sponge families with specimens from multiple collection sites (ANOVA, *p* > 0.05). Overall, Phloeodictyidae had the greatest evenness at the phylum level (Simpson evenness index value = 0.66), while the Farreidae microbiome was dominated by one archaeal phylum and had the lowest evenness (Simpson evenness index value = 0). Amongst families, the three Cladorhizidae sponges had the smallest range in Simpson evenness indices (mean ± SD, range) (0.55 ± 0.029, 0.055), while the two Euretidae had the greatest range (0.35 ± 0.194, 0.274).

**Figure 5.**
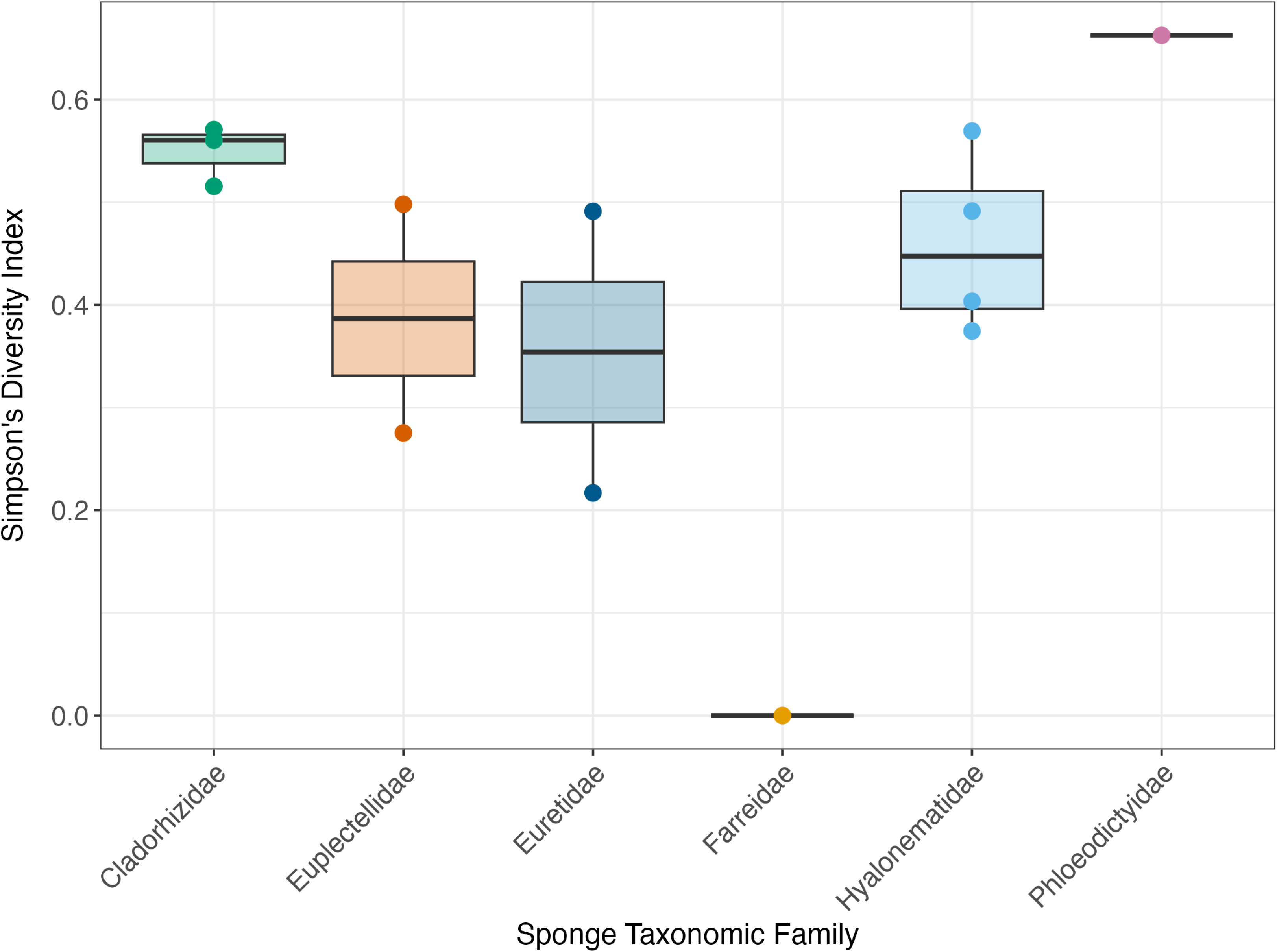
Microbial diversity across sponge taxa. Box plots of Simpson evenness values from sponge microbial communities. The center line of the box is the median, upper and lower bounds are quartiles. In order from left to right, sponge family Cladorhizidae, Euplectellidae, Euretidae, Farreidae, Hyalonematidae, and Phloeodictyidae.

Specimen Phloeodictyidae SH18796, likely an HMA sponge (Moitinho-Silva, Steinert, et al. 2017), had the greatest abundance and diversity of bacteria and archaea compared to the other sponges. In particular, it contained psychrophilic archaea *Thermoproteota*, *Cenarchaeum symbosium* (Preston et al. 1996).

The archaeal phylum *Thermoproteota* (mean ± SD, range) (59.7% ± 18.4%, 35.4%–93.5%) and the bacterial phylum *Proteobacteria* (78.2% ± 19.3%, 43.2%–99.3%) contributed the most to dissimilarity in microbiome composition between sponge families (SIMPER). These phyla together explained over 90% of the differences in most pairwise comparisons. The bacterial phylum *Chloroflexota* also contributed substantially to the microbiome composition of sponge family Phloeodictyidae, accounting for 34.3%–95.9% of community dissimilarity in pairwise comparisons with other Families.

### 3.4 Sponge Microbiome Metabolic Potential

Based on the presence of known metabolic genes, all sponge microbial communities had the capabilities to perform organic carbon oxidation, fermentation, and acetate oxidation (Figure 6A). Carbon degradation was overall driven by bacterial taxa. Carbon fixation genes (in addition to those in AOA) were present in microbiomes of all three Cladorhizidae, one Hyalonematidae, and the Phloeodictydiae. In particular, *Form 1 RuBisCO* was present in *Hyalonema* SH18898 and all three Cladorhizidae (SH18875, SH18725, and SH18747) and *Form 2 RuBisCO* was present in two of the Cladorhizidae (SH18725 and SH18747). Furthermore, the microbiome of Phloeodictyidae SH18796, contained genes related to methane cycling. Ten metagenome-assembled bacterial genomes of *Acidimicrobiales*, *Binatia*, *Chloroflexota*, *Dehaloccocoidia*, *Latescibacteria*, *Poribacteria*, and *Thermoanaerobacullia* contained methane monooxygenase regulatory protein B – *mmob.* In addition to *mmob*, MAG SH18796_Binatia_81_0 contained methane/ammonia monooxygenase subunits B and C – *pmoB, pmoC,* and C1 metabolism methanol oxidation – *mdh,* in sum constituting genetic potential for methyltrophy (Supplemental Table 4). Additionally, bacterial *Nitrospirales* MAG (SH18796_Nitrospirales_17_0) from Phloeodictyidae SH18796 was a nitrite-oxidizing autotroph containing genes for ATP citrate lyase (*aclA* and *aclB*) which enable bacterial autotrophy (Figure 6).

**Figure 6.**
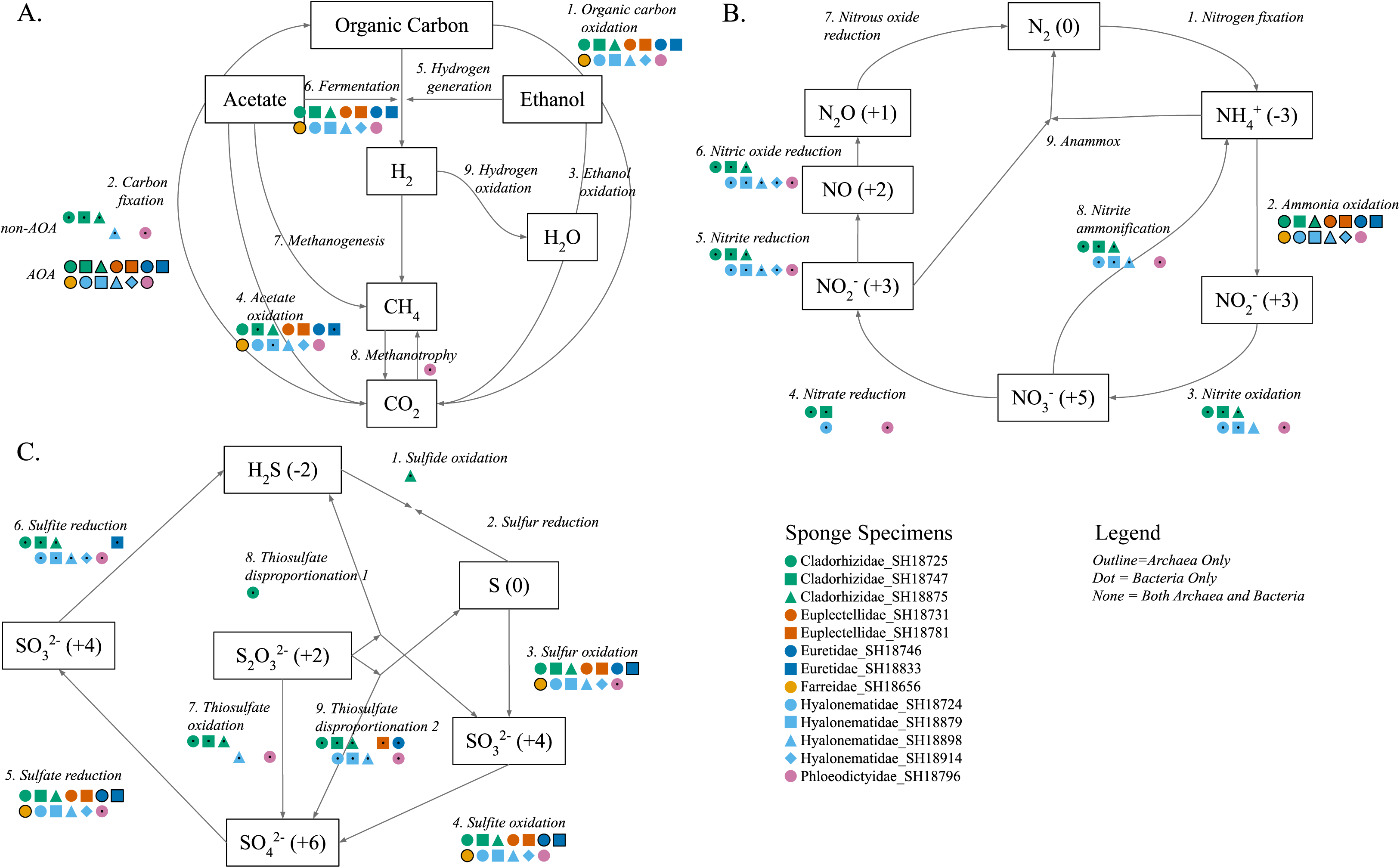
Metabolic potential for biogeochemical cycling in sponge-associated microbiomes. Functional genes involved in the carbon (A), nitrogen (B), and sulfur (C) cycles are shown as presence/absence for each sponge microbiome. Shape represents the sponge sample, color-coded by taxonomic family. If both Archaea and Bacteria contain genes to perform a given function, the shape is simply colored, while if it is only Bacteria there is a black dot in the middle of the shape, and an outline for only Archaea. (Cladorhizidae = Green, Euretidae = Dark Blue, Euplectellidae = Orange, Farreidae = Yellow, Hyalonematidae = Light Blue, Phloeodictyidae = Pink).

All sponge-associated microbial communities had the genetic capability to perform ammonia oxidation, due primarily to ammonia-oxidizing archaea (AOA) (*amoA, amoB, amoC).* Sponges Hyalonematidae SH18747 and Phloeodictyidae SH18796 also hosted bacteria that could perform ammonia oxidation (Figure 6B). No sponge microbiomes had the genetic capability to perform nitrogen fixation, nitrous oxide reduction, or annamox. All sponge microbiomes of Cladorhizidae, Hyalonematidae, and Phloeodictyidae contained bacteria that could perform nitrite reduction and nitric oxide reduction. Furthermore, all the Cladorhizidae, the Phloeodictyidae, and some of the Hyalonematidae contained bacteria that could perform nitrite oxidation (*nxrA, nxrB*), and nitrate reduction (*napA, napB*, *narG, narH*).

All sponge microbiomes had the genetic capabilities to perform sulfur oxidation, sulfite oxidation, and sulfate reduction, from bacterial and archaeal organisms (Figure 6C). Bacteria associated with a Cladorhidizae sponge (*Cladorhiza* SH18875) were capable of sulfide oxidation. Another Cladorhizidae sponge (*Chondrocladia* SH18725) also had bacteria capable of thiosulfate disproportionation (*phsA*), showing a distinct role for Cladorhizidae-associated microbes in sulfur cycling. Metabolic functions of sulfide oxidation, thiosulfate disproportionation (1 and 2), thiosulfate oxidation, and sulfite reduction were restricted to bacteria, not archaea, in the sponge microbiomes.

## 4.0 Discussion

### 4.1 Microbiome composition and structure across sponge taxa

The 13 siliceous sponges in this study were classified taxonomically into four Hexactinellida families and two Demospongiae families and harbored microbial communities characterized by relatively low overall diversity and a strong dominance of ammonia-oxidizing archaea (AOA). This lower microbial diversity and high AOA abundance in deep-sea sponge microbiomes has been previously reported (Busch et al. 2022a; Williams et al. 2024; Garritano et al. 2023) and contrasts with many shallow-water sponges, which often host greater diversities of bacteria and archaea across both high microbial abundance (HMA) and low microbial abundance (LMA) sponges (Thomas et al. 2016; Sabrina Pankey et al. 2022).

Our data and analyses showed that variation in microbiome composition across the 13 siliceous sponges was more closely associated with sponge taxonomy than collection site. Although sponge taxonomy was found to be a statistically significant driver of microbial community composition, limited sample sizes and loose clustering in ordination space (Figure 4) indicate that this relationship should be interpreted cautiously. Nevertheless, our results are consistent with previous research using operational taxonomic units (OTUs) of microbial communities associated with shallow sponges (Easson and Thacker 2014; Moitinho-Silva, Nielsen, et al. 2017; Leal et al. 2022), and deep marine sponges (Busch et al. 2022a; Kellner et al. 2018; Díez-Vives and Riesgo 2024; Reveillaud et al. 2014; Williams et al. 2024; Garritano et al. 2023) that similarly found that host taxonomy is a key driver of microbiome composition. Together, these findings suggest the possibility of a complex and long-standing co-evolutionary relationship between siliceous sponge hosts and their symbionts. Further sampling and sequencing of siliceous sponges is needed to further elucidate the relative contributions of host taxa, host traits such as feeding strategy and morphology, and environmental context on microbial community composition and function.

The distinct evolutionary history and lifestyles of the Demospongae specimens from Phloeodictyidae and Cladorhizidae in this study are reflected in their distinct microbial communities. Sponge Phloedictyidae SH18796 was the only HMA sponge (Moitinho-Silva, Steinert, et al. 2017) and thus microbiologically distinct compared to other LMA sponges in this study. In fact, when this sponge was included in the nMDS, it collapsed the heterogeneity between other sponge microbial communities (Supplemental Figure 1). Phloeodictyidae species consist of burrowing and non-burrowing sponges (Desqueyroux-Faúndez and Valentine 2002). Our specimen was a non-burrowing *Oceanapia* found on a rocky substrate, with a compact external crust and interior spreading fistules. Ecologically, this sponge had a similar niche to most other specimens in this study (i.e., as a sessile suspension feeder). Specimen Phloedictyidae SH18796 contained a notably higher relative abundance of bacterial phylum *Chloroflexota*, which contributed to 34.3%–95.9% of community dissimilarity in pairwise comparisons with other Families (SIMPER). Higher abundances of *Chloroflexota* have been previously reported in *Oceanapia* (Ruocco et al. 2021) and are consistent with characteristics typical in HMA sponges (Moitinho-Silva, Steinert, et al. 2017). The uniqueness of this sponge’s microbiome reinforces the importance of host taxonomy in shaping sponge microbiome composition.

Unlike other marine sponges which filter feed, Cladorhizidae sponges are carnivorous, capturing and digesting larger prey (e.g., small crustaceans) with hooks and enveloping prey with a thin membrane (Crew 2012; Vacelet and Boury-Esnault 1995). This distinct feeding behavior may potentially drive the conservation of the microbiome across the three Cladorhizidae included in this study — the three Cladorhizidae sponges had the smallest range in Simpson evenness amongst families. Notably, the two *Chondrocladia* specimens (SH18725 and SH18747) harbored highly similar microbial communities and contained bacterial phylum *Bacteriodota*: family *Flavobacteriaceae* in higher abundance than the other sponges. Previous researchers have reported higher abundances of *Flavobacteriaceae* in carnivorous sponges compared to other marine sponges (Verhoeven and Dufour 2018; Dupont et al. 2013), and our findings further suggest that *Flavobacteriaceae* may be a distinct part of the microbiome of Cladorhizidae, potentially related to carnivory. Similar to the Cladorhizidae, carnivorous plants also capture prey, and prey digestion in plants is facilitated by essential microbial symbionts (Sirová et al. 2018; Armitage 2017). Convergence in microbial structure and function in carnivorous pitcher plant microbiomes has been observed across continents (Bittleston et al. 2018). Cladorhizidae may experience similar selective pressures as terrestrial carnivorous plants, explaining in part the highly similar microbiomes of the three in this study.

The influence of sponge taxonomy on microbiome composition could be explained by vertical transmission of symbionts in siliceous sponges. All known sponge larvae are lecithotrophic, using energy provided maternally and not feeding during this life stage (S. Leys and Ereskovsky 2006). Studies on shallow Porifera have shown that larvae can contain microbial symbionts, confirming maternal transmission of sponge-associated microbes for at least some species (Schmitt et al. 2008; Maldonado 2007). It is unknown how widespread vertical transmission is among sponge taxa. Little is known about the life-cycle of siliceous sponges in the deep ocean, including how far larvae may disperse and their potential symbiont composition (reviewed in Carrier et al. 2022).

To better understand the role of host phylogeny in shaping the microbiome of siliceous sponge families, research comparing the microbiome of sponges from the same families across distinct deep ocean habitats is needed. In shallow sponges, it has been shown that within the same sponge taxa, strong changes in environmental conditions such as upwelling and proximity to anthropogenic pollution are associated with distinct microbiome structure (Batista et al. 2018). All the sponges in this study were from the vicinity of Puerto Rico, yet despite differences in site and depth (Supplemental Table 1), the microbial communities of the LMA sponges were relatively similar across samples (Figure 3). In some systems, the microbiome is known to facilitate host adaptation to changing environments (Baldassarre et al. 2022; Ziegler et al. 2017; Petersen et al. 2023), and it remains unknown if and how the microbiome of deep-sea siliceous sponges may potentially allow for host adaptation to different environments.

### 4.2 Bacterial Community Composition

All sponges except the Farreidae SH18656 contained bacterial phylum *Proteobacteria*. The most dominant classes of *Proteobacteria* across all sponges were *Gammaproteobacteria* and *Alphaproteobacteria*. The presence and abundance of *Thermoproteota* and *Proteobacteria* suggests that these groups may play fundamental roles in siliceous sponge microbiomes and compose a “core microbiome” (Neu et al. 2021; Shade and Handelsman 2012).

Three of the four Hyalonematidae sponges (SH18724, SH18879, and SH18898) shared higher abundances of bacterial phylum *Nitrospinota*, unique from the other groups of sponges. In addition to belonging to a different subclass of Hexactinellida, the Hyalonematidae are also morphologically distinct from other sponges in this dataset – they have long anchoring spicules (basalia), which act as stalks suspending the body of the sponge above the seafloor. Current velocity and turbulence vary across the benthic boundary layer, so organisms that are elevated above the seafloor experience different physical and biogeochemical conditions than those lower down (Vogel 1996). Perhaps the Hyalonematidae’s height in the benthic boundary layer exposes them to different fluid shear currents and nutrient flux, enabling greater abundance of these nitrite oxidizers. Within Hexactinellida, order Amphidiscosida, basalia length varies greatly, and further sampling is required to investigate this potential trend.

Although host phylogeny appears to play a role in shaping microbiome structure, the microbial communities also contain pathogens and symbionts that may be transient. Two of the sponges (Euplectellidae SH18781 and Cladorhizidae SH18875) contained *Patescibacteria*, a bacterial phylum from the Candidate Phyla Radiation, and Euplectella SH18731 was the only sponge to contain *Bdellovibrionata*. *Patescibacteria* are ultrasmall and have symbiotic and parasitic relationships with bacterial hosts (Brown et al. 2015; Castelle et al. 2018). Similarly, *Bdellovibrionata* are often parasitic on gram-negative bacteria (Shilo and Bruff 1965; Jurkevitch et al. 2000). The presence of these bacteria with potentially parasitic and symbiotic lifestyles suggest that these microbiomes are dynamic, which warrants further investigation and sampling to establish a “core” versus transient siliceous sponge-associated microbial community.

### 4.3 Ammonia-oxidizing archaea dominate siliceous sponge microbiome

The microbial communities of the siliceous sponges were dominated by ammonia-oxidizing archaea (AOA) from the family *Nitrosopumilacaea.* Particularly striking was that the two strains of *Nitrosopumilaceae* exclusively composed the microbiome of Farreidae SH18656. To our knowledge, such extreme dominance by a single archaeal taxon has rarely been reported in sponge-associated microbiomes and suggests strong host filtering or specialization.

AOA are autotrophic and could provide a carbon source for the sponge host in the deep ocean, as photosynthetic symbionts provide carbon sources for shallow sponges in low light environments (Hudspith et al. 2022b). Furthermore, the role of AOA in oxidizing ammonia, a byproduct of sponge metabolism, may be crucial in maintaining sponge host health, as high abundances of ammonia can be toxic to aquatic organisms (reviewed in Camargo and Alonso 2006). Thus, the high abundance of AOA could be a key factor in the ecology and biogeochemical function of these deep-sea sponges (Glasl et al. 2024). Previous research has shown that *Nitrosopumilacaea* species phylogeny is closely related to sponge host taxonomy (Garritano et al. 2023). This close relationship between *Nitrosopumilacaea* and host sponge phylogeny and the high abundance of *Nitrosopumilacaea* found here suggests that these archaea and siliceous sponges may have a complex co-evolutionary history.

Interestingly, Hyalonematidae SH18879 did not contain a high abundance of AOA, but contained *Nitrospinota* (nitrite-oxidizing bacteria) in higher abundance than other sponges, potentially compensating for the lower AOA. *Nitrospinota* have been detected in other deep-sea sponge microbiomes (Reveillaud et al. 2014; Moitinho-Silva, Nielsen, et al. 2017), where they are typically present in low abundance and contribute to nitrification alongside AOA. However, most studies on both shallow and deep-sea sponges report consistently high abundances of AOA (Bayer et al. 2020; Garritano et al. 2023; Moeller et al. 2019), suggesting that the relative depletion of AOA and enrichment of *Nitrospinota* may reflect a distinct host-symbiont association or localized environmental adaption.

In shallow sponges, AOA are present in the larval microbiome, suggesting their importance to the ontogeny of marine sponges potentially through nitric oxide signaling, a chemical regulator of cellular function (Glasl et al. 2024; Moeller et al. 2019; Schläppy et al. 2010). The microbiome is known to modulate the timing of life history transitions in hosts; for example in *Drosophila* development (Shin et al. 2011), mosquito metamorphosis (Coon et al. 2014), and plant flowering (Wagner et al. 2014). It remains unknown why one species would rely on another species for such critical aspects of organism fitness across the tree of life (reviewed in Koskella et al. 2017). Future investigation of AOA in siliceous sponges and their larvae may shed further light on the natural selection processes acting on host, microbes, and host-microbe interactions.

### 4.4 Siliceous sponge microbiome plays important role in nutrient cycling

The microbial communities in siliceous sponges play a vital role in cycling nutrients in the nutrient-poor oligotrophic deep ocean, referred to as the “sponge-loop” (Busch et al. 2022a; de Goeij et al. 2013). Our results show that the microbial communities associated with the 13 sponges have the genetic capabilities to perform key biogeochemical functions, particularly in the cycling of carbon, nitrogen, and sulfur. Deep-sea sponges are heterotrophic, and thus the function of the microbiome involves breaking down of organic matter for the acquisition of carbon and nutrients (Busch et al. 2022a). For each individual sponge, its capacity to influence nutrient cycling (carbon, nitrogen, and sulfur) varied depending on the predicted metabolic function of the MAGs (Supplemental Table 4).

In regard to carbon cycling, all sponge-associated microbial communities contained genes encoding for organic carbon oxidation, fermentation, and acetate oxidation (Figure 6A), supporting microbial and host sponge carbon metabolism. These presence of the pathways show microbial potential to break down organic matter (DOM and POM) into simpler compounds, producing energy and metabolites which may be 1) directly utilized by the microbiome and/or sponge host for growth and 2) released into the surrounding seawater, indirectly supporting deep-sea communities as has been show in shallow sponges (de Goeij et al. 2013; Rix et al. 2017).

While sponge microbiomes contained a wide range of sulfur-metabolism capabilities (Figure 6C), broad metabolic potential was found in the Cladorhizidae specimens. In particular, one of the Cladorhizidae in this study (*Cladorhiza* SH18875) contained genes for the removal of sulfide. These findings are also consistent with the hypothesis that sulfur-cycling bacterial symbionts have also evolved close associations with sponge hosts, similar to AOA (Ramírez et al. 2023).

In addition to the high abundance of AOA and the importance of nitrification in the siliceous sponge microbiomes, there were also bacteria with capabilities to reduce nitrite, nitrate, and nitric oxide (Figure 6B). Dissimilatory nitrate reduction to ammonium (DNRA) was also present, which produces ammonia, providing another source of ammonia substrate likely metabolized by the high abundance of AOA. This microbial cross feeding and nutrient competition in the nitrogen cycle may potentially shape the microbial community composition; previously researchers showed that metabolic dependencies may enable microbial species co-occurrence (Zelezniak et al. 2015).

### 4.5 Evidence of Potential for Microbial Autotrophy

In addition to the chemolithoautotrophic AOA, we found evidence for autotrophy in the microbial communities associated with six of the siliceous sponges. Previous research on the microbiomes of shallow sponges have shown that they sometimes contain photosynthetic symbionts which can contribute to host sponge metabolism and ecosystem primary productivity (Clive R. Wilkinson 1987; C. R. Wilkinson 1983; Hudspith et al. 2022a).

Bacteria associated with one *Hyalonema* and all three Cladorhizidae sponges contained *RuBisCO*, an enzyme responsible for the majority of photosynthetic carbon fixation on Earth (Geider et al. 2001; Bar-On and Milo 2019; Raven 2009). Specifically, of the three forms of RuBisCO; *Form 1 RuBisCO* was present in the *Hyalonema* and all three Cladorhizidae and *Form 2 RuBisCO* was present in the two of the Cladorhizidae: *Chondrocladia*. Recent research has suggested that RuBisCO may play a large role in the fixation of carbon in the dark ocean (Jaffe et al. 2025). Due to their carnivorous feeding mechanisms, Cladorhizidae sponges may benefit from microbial autotrophy when prey is not available to feed heterotrophic metabolism. The presence of RuBisCO genes in the microbiome of these sponges suggests an adaptive mechanism for nutrient acquisition in low-resource environments. Previously, researchers reported evidence of genetic potential for microbial autotrophy associated with deep-sea Cladorhizidae sponges via methanotrophy, but not via RuBisCO, in the open ocean (Hestetun et al. 2016) and near methane seeps (Jean et al. 1996).

Absent in the Cladorhizidae in this study, the microbiome of sponge Phloeodictyidae SH18796 included ten MAGs from a range of bacterial taxa containing methane monooxygenase regulatory protein B – *mmob.* This may confer methane oxidation potential in these organisms, but unlikely full methanotrophy in the absence of additional genes. However, one organism, SH18796_Binatia_81_0, had potential for methyltrophy containing *mmob* in addition to methane/ammonia monooxygenase subunits B and C – *pmoB* and *pmoC*, and C1 metabolism methanol oxidation – *mdh*. Methyltrophy potential in this organism is consistent with recent research on the taxonomic group (Murphy et al. 2021) (previously *Candidatus* Phylum UBP10, presently class *Binatia*). The presence of genes involved in methyltrophy was surprising considering the absence of methanogens or a methane seep near our study sites. Bacteria capable of methanotrophy have been reported in shallow sponges without a clear methane source (Ramírez et al. 2023). Incomplete oxidation of methane in sediment may potentially allow methane to diffuse into the seawater, thus providing a methane source for these metabolisms.

Genes encoding ATP citrate lyase, a key enzyme in the reverse TCA cycle, were present in nitrite-oxidizing bacteria associated with specimen Phloeodictyidae SH18796. Because this pathway is used by chemolithoautotrophic microbes to fix carbon (reviewed in Hügler and Sievert 2011), its presence suggests that sponge-associated microbes may supplement sponge heterotrophy by providing an additional carbon source.

## 5.0 Conclusion

Deep-sea siliceous sponges are long-lived and ecologically important members of marine ecosystems, yet their microbial symbioses remain among the least studied within Porifera. Here, metagenomic sequencing revealed microbial community composition and function associated with siliceous sponges from five taxonomic families. Sponge-associated microbial communities were influenced by sponge host phylogeny and had the genetic potential to contribute to carbon, nitrogen, and sulfur cycling, spanning both heterotrophic and autotrophic metabolisms, with ammonia-oxidizing archaea dominating most specimens. Future sampling, sequencing, and microscopy across environmental gradients and sponge life stages may further clarify the dynamics between siliceous sponges and their symbionts, such as establishment, persistence, and spatial organization. As the ocean changes and benthic ecosystems respond and adapt, establishing genome-resolved baselines for siliceous sponge microbial symbioses is critical to evaluating ecosystem health. Together, the results of this research highlight the important role of microbial communities in siliceous sponge ecology and deep-sea nutrient cycling, furthering our understanding of these enigmatic organisms.

## Data Availability

Raw sequencing reads and metagenome-assembled-genomes (MAGs) are publicly available from National Center for Biotechnology Information (NCBI) BioProject Accession: PRJNA1355197. Supplemental tables contain metadata associated with each sponge (Supplemental Table 1), sequencing and assembly statistics (Supplemental Table 2), statistics about each MAG (Supplemental Table 2), functional annotations of MAGs (Supplemental Table 4), and phylogenetic tree associated with Figure 3 (Supplemental File 1). Code used for data analysis is available on Github: (https://github.com/kate-lane/siliceous-sponge-microbes).

## Acknowledgements

We thank the captain, crew, and science team of the *R/V Falkor (too)* and the pilots and engineers of ROV *SuBastian* for their expertise and support during sample collection on cruise FKt230417. Ship time was provided by the Schmidt Ocean Institute (to C.M.H.). We are grateful to the Smithsonian National Museum of Natural History for their curatorial support, taxonomic identification, and housing of specimens. Genetic sequencing was performed at the Roy J. Carver Biotechnology Center at the University of Illinois at Urbana-Champaign and we thank Alvaro Hernandez and staff for support. Computational analyses were conducted using the Poseidon High Performance Computing Cluster at Woods Hole Oceanographic Institution and we thank the technical support staff, including Gretchen Zwart. This work was supported by the Woods Hole Oceanographic Institution – Massachusetts Institute of Technology Joint Program Ocean Ventures Fund (to K.R.L.) and the American Academy of Underwater Sciences Ruth B. Turner Scholarship (to K.R.L.). K.R.L was supported by the National Science Foundation Graduate Research Fellowship (DGE1252522).

## Figure Legends

**Supplementary Figure 1.** nMDS ordination of microbial community composition across siliceous sponges including Phloeodictyidae. Non-metric multidimensional scaling plot (nMDS) based on Bray Curtis similarities of sponge microbiome phyla composition and abundance. Closely clustered symbols correspond to samples with similar Bacteria and Archaeal phyla composition and abundance. Sponge family taxa are indicated by color. Locations of specimens are indicated by shape (Desecheo Ridge = Square, Esperanza Ridge = Triangle, Ridge SW of Vieques = Circle, Whiting Seamount = Diamond.) Letters adjacent to symbols indicate individual specimens as labelled in Figure 1: Sponge Plate. The distinctness of the Phloeodictyidae sponge compared to the others in this dataset collapses the differences between other specimens.

